# Gut microbiome is associated with recurrence-free survival in patients with resected Stage IIIB-D or Stage IV melanoma treated with immune checkpoint inhibitors

**DOI:** 10.1101/2024.04.16.589761

**Authors:** Mykhaylo Usyk, Richard B. Hayes, Rob Knight, Antonio Gonzalez, Huilin Li, Iman Osman, Jeffrey S. Weber, Jiyoung Ahn

**Affiliations:** Department of Population Health, NYU Grossman School of Medicine, New York, NY, USA; NYU Laura and Isaac Perlmutter Cancer Center, New York, NY, USA; Departments of Pediatrics, Computer Science & Engineering, and Bioengineering; Center for Microbiome Innovation, University of California, San Diego, La Jolla, CA; The Ronald O. Perelman Department of Dermatology, NYU Grossman School of Medicine, New York, NY, USA; Department of Medicine, NYU Grossman School of Medicine, New York, NY, USA

## Abstract

The gut microbiome (GMB) has been associated with outcomes of immune checkpoint blockade therapy in melanoma, but there is limited consensus on the specific taxa involved, particularly across different geographic regions. We analyzed pre-treatment stool samples from 674 melanoma patients participating in a phase-III trial of adjuvant nivolumab plus ipilimumab versus nivolumab, across three continents and five regions. Longitudinal analysis revealed that GMB was largely unchanged following treatment, offering promise for lasting GMB-based interventions. In region-specific and cross-region meta-analyses, we identified pre-treatment taxonomic markers associated with recurrence, including *Eubacterium, Ruminococcus, Firmicutes*, and *Clostridium*. Recurrence prediction by these markers was best achieved across regions by matching participants on GMB compositional similarity between the intra-regional discovery and external validation sets. AUCs for prediction ranged from 0.83-0.94 (depending on the initial discovery region) for patients closely matched on GMB composition (e.g., JSD ≤0.11). This evidence indicates that taxonomic markers for prediction of recurrence are generalizable across regions, for individuals of similar GMB composition.

**Highlights:**

- Overall gut microbiome (GMB) composition is largely unchanged during ICB treatment.
- GMB composition varies by geographic region
- We identified gut bacterial markers associated with recurrence in region-specific analyses.
- Region-identified markers are generalizable if GMB composition is taken into account by matching.

## Introduction

Melanoma is the 6^th^ most common form of cancer in the U.S., accounting for approximately 100,000 new cases annually (Siegel et al., 2023). Immune checkpoint blockade (ICB), utilizing monoclonal antibodies targeting programmed death 1 (PD-1) and cytotoxic T-lymphocyte antigen 4 (CTLA-4), are treatment options that can provide durable benefit in metastatic and high-risk resected melanoma. However the benefit of ICB is unpredictable and 25-30% of those treated experience cancer recurrence (Weber et al., 2023). Identifying robust biomarkers to predict treatment outcomes is imperative. Predictive markers may support personalized treatment plans, resulting in improved patient management, ultimately enhancing treatment efficacy and outcomes.

Accumulating evidence suggests that the gut microbiome (GMB) influences survival, progression and recurrence in ICB-treated melanoma (Gopalakrishnan et al., 2018; Matson et al., 2018; Routy et al., 2018; Peters et al., 2019). These associations have been further supported by intervention experiments, in clinical trials and animal models, which have demonstrated the potential for improved outcomes in melanoma through fecal microbiome transplant (FMT) (Baruch et al., 2021; Davar et al., 2021; McQuade et al., 2020). Notably, clinical trials conducted by Davar *et al*. (Davar et al., 2021) and Baruch *et al*. (Baruch et al., 2021) showed evidence of ICB response in treatment-refractory patients following FMT, associated with consistent activation of CD8+ T cells. Additionally, several pre-clinical human-to-mouse FMT transplant studies demonstrated similar T-cell activity in anti-PD-L1-based therapies for melanoma (Gopalakrishnan et al., 2018; Matson et al., 2018). Furthermore, studies showed that a high-fiber diet in melanoma patients undergoing ICB in the neoadjuvant setting was related to alteration in GMB and enhanced treatment response (Simpson et al., 2022), with further confirmation in a pre-clinical mouse model where high-fiber treatment was associated with changes in the GMB and improved treatment outcomes (Spencer et al., 2021).

Although these studies provide promising clinical insights, the identified bacterial markers for predicting treatment outcomes in melanoma have varied considerably among studies (Lee et al., 2022). In fact, in a recent multi-regional analysis of European patients, Lee *et al* argued that GMB markers are region specific (Lee et al., 2022). While this discrepancy in bacterial marker identification, by region, may be attributed to clinical selection criteria, different ICB treatment modalities, small sample sizes, or population-specific characteristics (He et al., 2018b; Lee et al., 2022), it is becoming evident that geographic variation in compositional attributes likely plays an important role (Lee et al., 2022), as geography, as well as relocation, are known to be strong determinants of GMB composition (He et al., 2018b; Kaplan et al., 2019; Peters et al., 2020; Vangay et al., 2018). This underscores the critical need to sample the microbiome from diverse patient groups and geographic areas to comprehensively capture GMB biodiversity and identify robust bacterial markers for treatment outcomes and their associated contexts.

For the current investigation, we studied participants in the Checkmate 915 randomized, double-blind, phase III trial (ID: NCT02388906), that was composed of 1,833 patients who received nivolumab 240 mg once every 2 weeks plus ipilimumab 1 mg/kg once every 6 weeks (916 patients) or nivolumab 480 mg once every 4 weeks (917 patients) for ≤ 1 year. In this cohort, we investigated the association between GMB and melanoma recurrence in 674 trial participants who provided stool samples. This study was carried out under a standard protocol across five broad geographic regions, allowing for a detailed analysis of the regional association of GMB with treatment outcomes and allowing us to directly address the issue of geographic variation while maximizing bias control via rigorous clinical trial design. In pre-treatment stool samples using shotgun metagenomics, we achieved strain-level resolution of the GMB. We demonstrate broad generalizability of certain strains of bacteria in meta-analysis and more robust cross-regional prediction, overcoming previous replication hurdles, via GMB matching. Additionally, we sequenced stool samples collected at 7 weeks and 29 weeks after treatment initiation in a sub-sample, to assess the stability of the GMB following ICB treatment. This investigation is the first to explore the GMB in melanoma patients in the adjuvant setting, potentially uncovering crucial insights that could lead to more effective, personalized treatment strategies to improve patient outcomes.

## Results

### Patient Characteristics

Our prospective study of GMB and melanoma included 674 patients with resected stage IIIB-D or IV melanoma, who were randomized to receive adjuvant nivolumab plus ipilimumab or nivolumab alone (Weber et al., 2023). All participants provided a stool sample prior to ICB treatment and approximately half of the patients provided stool samples post-treatment (at weeks 7 and 29 follow-up visits) (**Supplemental Figure 1**). The 674 patients were evenly distributed between the combination therapy and nivolumab monotherapy arms (**Table 1**). Patients were majority white (99.0%) and male (58.9%), and the mean (SD) age was 55.0 (13.9) years. Melanoma recurrence was similar for the combination (35.0%) and nivolumab arms (39.8%), similar to what was previously reported for the full trial series (35.4% vs. 36.8%)(Weber et al., 2023).

**Table 1.** Demographic and Clinical Characteristics of Study Participants, Stratified by Geographic Region.

| Variable | Overall | North America* | Australia | Western Europe | Eastern Europe | ROW** |
| --- | --- | --- | --- | --- | --- | --- |
| N | 674 | 95 | 127 | 367 | 43 | 42 |
| Age Mean ± SD | 55.03 ± 13.88 | 55.29 ± 14.26 | 55.31 ± 13.73 | 54.72 ± 13.73 | 51.77 ± 15.17 | 58.9 ± 12.68 |
| Gender |  |  |  |  |  |  |
| Female | 277 (41.1%) | 40 (42.1%) | 44 (34.6%) | 160 (43.6%) | 19 (44.2%) | 14 (33.3%) |
| Male | 397 (58.9%) | 55 (57.9%) | 83 (65.4%) | 207 (56.4%) | 24 (55.8%) | 28 (66.7%) |
| Race |  |  |  |  |  |  |
| White | 667 (99.0) | 91 (95.8%) | 126 (99.2%) | 366 (99.7%) | 43 (100%) | 41 (97.6%) |
| Other | 7 (1.0%) | 4 (4.2%) | 1 (0.8%) | 1 (0.3%) | 0 (0%) | 1 (2.4%) |
| Recurrence |  |  |  |  |  |  |
| Yes | 252 (37.4%) | 32 (33.7%) | 49 (38.6%) | 137 (37.3%) | 24 (55.8%) | 10 (23.8%) |
| No | 422 (67.6%) | 63 (66.3%) | 78 (61.4%) | 230 (62.7%) | 19 (44.2%) | 32 (76.2%) |
| Median Follow-up Time (months) ± IQR | 24.9 ± 19.8 | 23.4 ± 17.5 | 24.9 ± 22.1 | 24.8 ± 19.7 | 15.0 ± 25.7 | 27.6 ± 3.0 |
| Trial Arm |  |  |  |  |  |  |
| Nivo240mg+Ipi1mg/kg | 346 (51.3%) | 45 (47.4%) | 62 (48.8%) | 190 (51.8%) | 25 (58.1%) | 24 (57.1%) |
| Nivolumab 480mg | 328 (48.7%) | 50 (52.6%) | 65 (51.2%) | 177 (48.2%) | 18 (41.9%) | 18 (42.9%) |
| Stage |  |  |  |  |  |  |
| IIIB | 201 (29.8%) | 29 (30.5%) | 39 (30.7%) | 113 (30.8%) | 11 (25.6%) | 9 (21.4%) |
| IIIC | 366 (54.3%) | 57 (60%) | 62 (48.8%) | 199 (54.2%) | 26 (60.5%) | 22 (52.4%) |
| IIID | 15 (2.2%) | 4 (4.2%) | 1 (0.8%) | 9 (2.5%) | 0 (0%) | 1 (2.4%) |
| IV | 92 (13.7%) | 5 (5.3%) | 25 (19.7%) | 46 (12.5%) | 6 (14%) | 10 (23.8%) |
| BRAF Mutation |  |  |  |  |  |  |
| Wildtype | 325 (48.2%) | 47 (49.5%) | 67 (52.8%) | 182 (49.6%) | 19 (44.2%) | 10 (23.8%) |
| Mutant | 188 (27.9%) | 20 (21.1%) | 35 (27.6%) | 111 (30.2%) | 19 (44.2%) | 3 (7.1%) |
| Missing | 161 (23.9%) | 28 (29.5%) | 25 (19.7%) | 74 (20.2%) | 5 (11.6%) | 29 (69%) |
| LD*** (L/U, mean ± SD) | 216.6 ± 86.5 | 186.8 ± 84.3 | 189.7 ± 3 | 230.5 ± 93.3 | 187.4 ± 52.3 | 276.8 ± 109.5 |
| Missing (N) | 12 | 0 | 1 | 9 | 1 | 1 |

\* North America (NA: U.S. and Canada); \*\* ROW stands for “rest of world”, patients from Brazil (n = 28) and New Zealand (n = 14).
\*\*\* LD refers lactate dehydrogenase (LDH) level and L/U represent the Lower/Upper of the test result

Global beta diversity analysis using Jensen-Shannon divergence (JSD, a measure of GMB similarity between pairs of samples) revealed that GMB differed significantly by region, sex, stage, and gender, both when performing univariate and co-adjusted analyses (**Figure 1A**). The gut microbiome compositions from North American (USA and Canada) and Eastern European participants showed the greatest pairwise dissimilarity (R^2^=4.84%, p=0.001), while those from Eastern and Western European participants, the two most proximal areas, showed minimal differences (PERMANOVA R^2^ = 0.52%, p-value=0.034 (**Figure 1B**).

**Figure 1.**
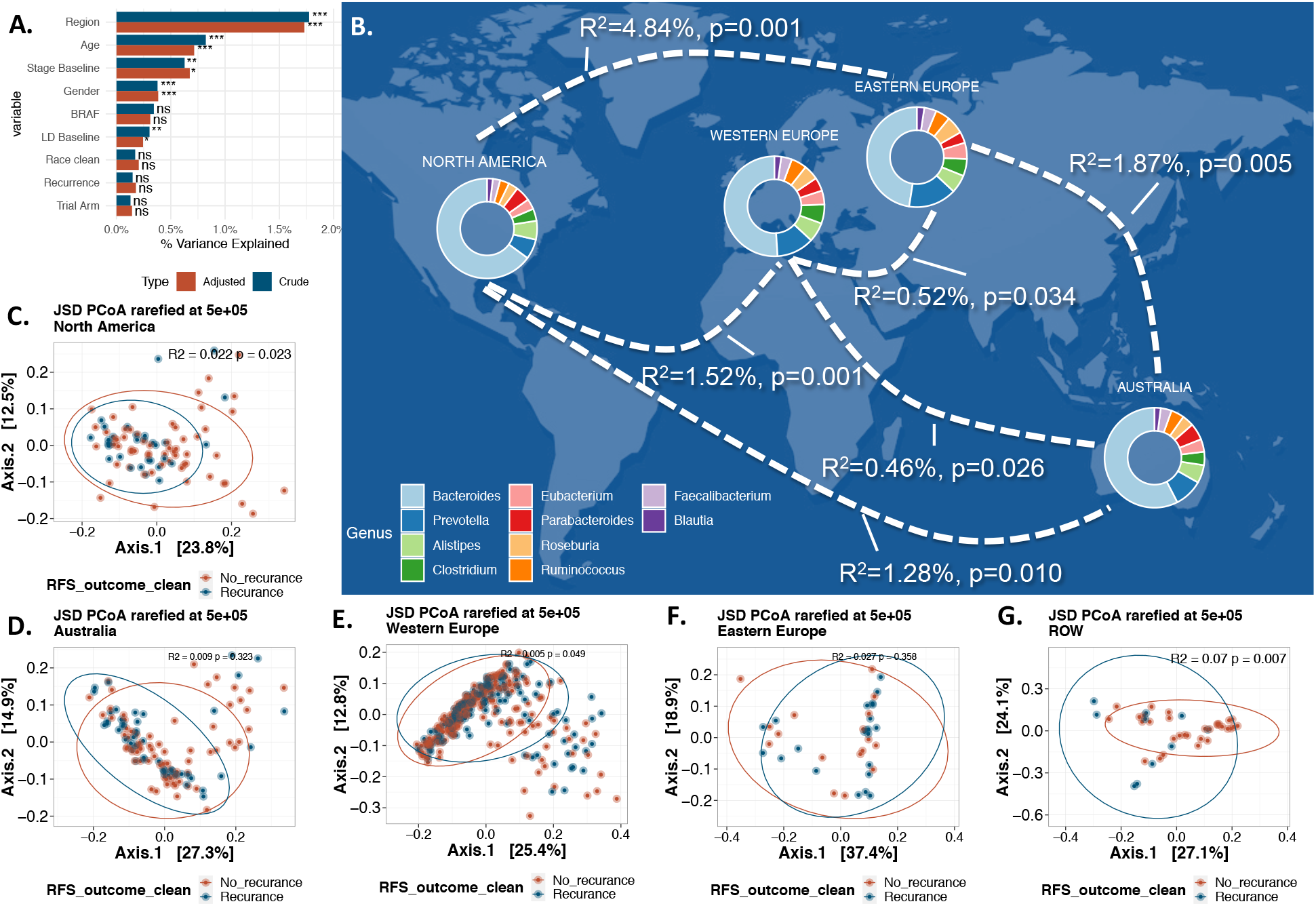
Beta Diversity, Regional Variation and Melanoma Recurrence. (A) illustrates a PERMANOVA analysis of essential clinical and demographic variables within our study, using JSD distance. Color indicates the analysis type: crude in blue and adjusted (adjustment for each other variable) in orange. The x-axis denotes R^2^, reflecting the proportion of overall gut microbiota composition variance, with stars adjacent to the bars indicating significance (p-values: 0.05 *, 0.01 **, 0.001 ***, >0.05 NS). Panel (B) presents a map of the geographic regions, with paired PERMANOVA results for each geographic pair displayed on the plotted curves. All pairs exhibited significant differences. Donut charts plotted over each geographic region represent the top 10 genera across (based on abundance) all samples within that region. Panels (C-G) depict the principal coordinate analysis (PCoA) for each of the five geographic regions, considering recurrence as the outcome (R^2^ and p-values are provided for each region).

### GMB and Recurrence

GMB structure (beta diversity) was not associated with melanoma recurrence in the overall study of 674 patients (in both crude and adjusted analysis, **Figure 1A**). These relationships were similar for the two arms of the trial (R^2^=0.003, p-value=0.67 and R^2^=0.003, p-value=0.38, PERMANOVA for combination and mono treatment respectively). In region-stratified analysis, GMB beta diversity was associated with recurrence in North America (R^2^=0.022, p-value=0.023), Western Europe (R^2^=0.005, p-value=0.049) and rest of world (Brazil and New Zealand) (ROW) (R^2^=0.07, p-value=0.007); there was no evidence of association for Australia (R^2^=0.009, p-value=0.32) or Eastern Europe (R^2^=0.027, p-value=0.36) (**Figure 1C-G**). Because of the differential GMB associations by geographic regions, subsequent analyses are based on region-specific analyses.

In region stratified analyses using ANCOM-BC, we identified several GMB taxa associated with recurrence. Nine bacterial taxa were associated with recurrence in North America (**Figure 2A** – dark green points and **Supplemental Table 1**): *Eubacterium sp. CAG:115, Ruminococcus sp. CAG:177, Eubacterium sp. CAG:786, Eubacterium siraeum, Firmicutes bacterium CAG:137, Clostridium sp. CAG:780, Clostridiales bacterium 1-7-47, Firmicutes bacterium CAG:884, Aeromonas salmonicida,* and *Peptostreptococcus anaerobius*. Among these, bacteria belonging to the genera of *Eubacterium, Ruminococcus, Firmicutes*, and *Clostridium* have previously been identified as predictive of recurrence in melanoma (Lee et al., 2022; Matson et al., 2018; Routy et al., 2018), while *Aeromonas salmonicida* and *Peptostreptococcus anaerobius* represent novel markers. In Western Europe (**Figure 2A** – brown points), two novel markers, *Bariatricus massiliensis* and *Blautia schinkii,* were identified. In ROW we identified a Clostridium, *Clostridiales bacterium* 1-7-47FAA, while for Eastern Europe, *Lawsonia intracellularis*, a novel marker, was the only recurrence-associated taxa identified.

**Figure 2.**
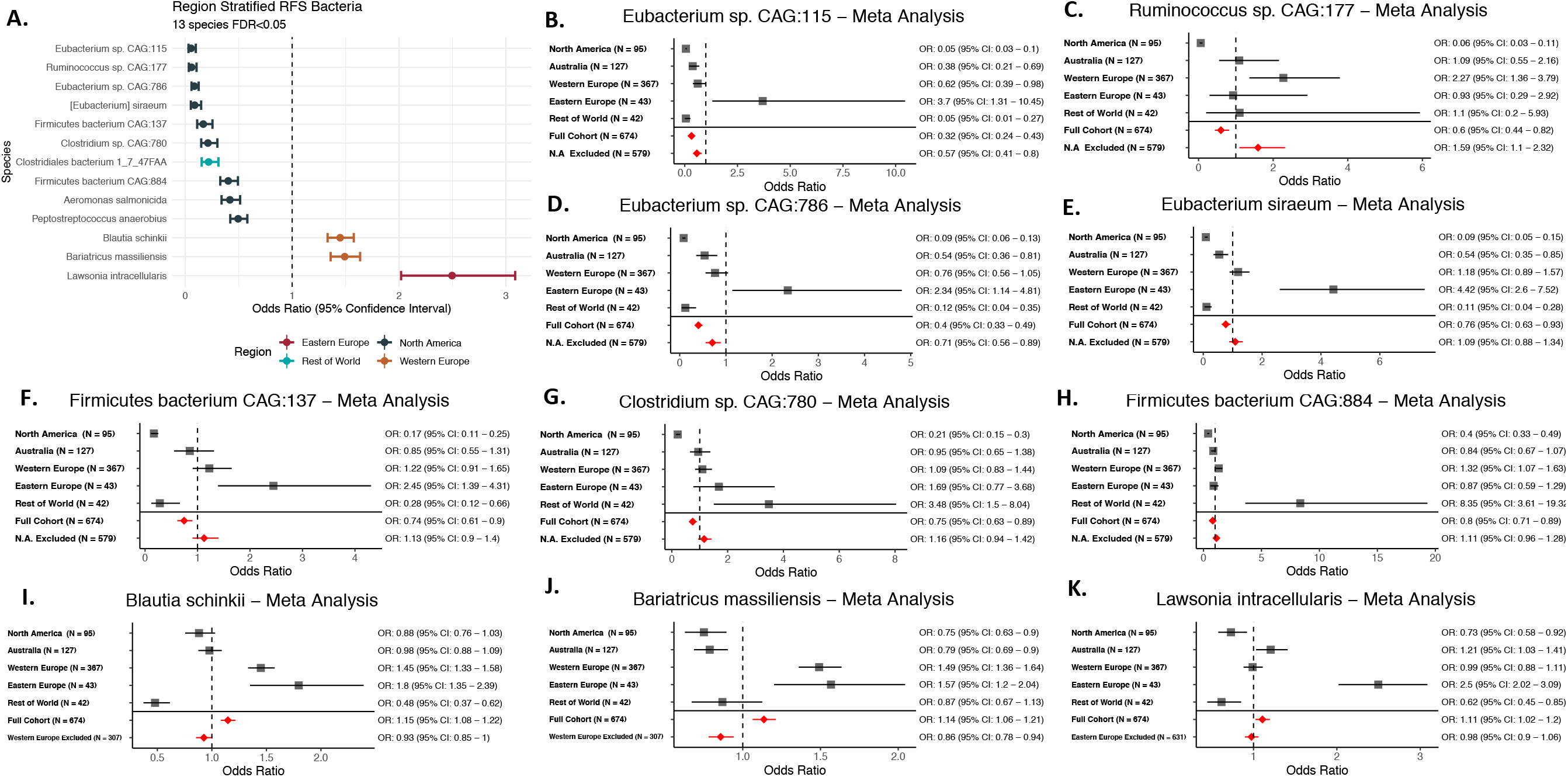
Region Stratified Analysis of Individual GMB Taxa and Melanoma Recurrence. (A) shows the region-stratified analysis as a forest plot with each point and associated confidence interval colored by the geographic region in which the identified gut microbiome markers prospectively associated with melanoma recurrence. All strains shown are significant after adjustment for multiple testing (FDR<0.05), with effects adjusted for participant age, sex, tumor stage, BRAF mutation and study arm. (B-K) show the meta-analysis of region-specific gut microbiome markers associated with recurrence in melanoma patients across geographic regions. Each panel shows analysis of a specific microbiome strain by region and, in meta-analyses, for the full cohort and for the full cohort minus the discovery region in (A) (N.A. stands for North America in the exclusion line). Meta analyses were performed using random-effect meta-analysis models (Schwarzer, 2007). Bacteria from (A) that are not significant in meta-analysis (*Aeromonas salmonicida, Clostridiales bacterium 1-7-47 FAA* and *Peptostreptococcus anaerobius*) are not shown in (B-K).

We performed a meta-analysis on these region-specific markers across all regions, to determine whether the markers associated with recurrence in one region were generalizable. Although there is significant heterogeneity between regions, we found that seven regionally identified recurrence markers were also associated with recurrence in cross-region meta-analyses (**Figure 2B-K)**, including markers initially identified for North America (*Eubacterium sp. CAG:115, Ruminococcus sp. CAG:177, Eubacterium sp. CAG:786, Eubacterium siraeum*), Western Europe (*Bariatricus massiliensis* and *Blautia schinkii*), and Eastern Europe (*Lawsonia intracellularis*). In meta-analyses excluding the original discovery region, *Eubacterium sp*.

*CAG:115* and *Eubacterium sp. CAG:786* remained significantly associated in the same (protective) direction, indicating the potential role of these bacteria in a general context, while *Ruminococcus sp. CAG:177, Bariatricus massiliensis* and *Blautia schinkii* remained significant, but showed an inverse association (**Figure 2B-K**, bottom common effect shown in red). This suggested a potential GMB context specificity of taxonomic markers for recurrence; that is, specific GMB markers may predict recurrence given a specific GMB composition. This is explored in “*GMB matching facilitates cross-regional generalizability*” section below.

To explore the potential functional mechanisms of microbial association with recurrence, we investigated the association between recurrence-associated taxa, from region-stratified analyses (see **Figure 2A**), and KEGG Level 3 pathways. We identified 57 functional pathways linked to recurrence-associated species (FDR <0.0001) (**Figure 3A**). Fifty-five of the 57 pathways were classified as “Metabolism” at KEGG Level 1, with the remaining two are involved in the biosynthesis of secondary metabolites. Given prior findings illustrating a connection between fiber consumption, gut microbiota shifts, and improved melanoma outcomes during treatment (Spencer et al., 2021), we focused on 15 carbohydrate-associated “Metabolism” pathways with correlations >0.3, including for amino-sugar and nucleotide-sugar metabolism (**Figure 3B**). Of the 15 carbohydrate-associated pathways, 8 were differentially associated with recurrence in the North American region, but not in other regions (FDR<0.0001, adjusted for age, sex, tumor stage, BRAF mutation and study arm) (**Figure 3C**).

**Figure 3.**
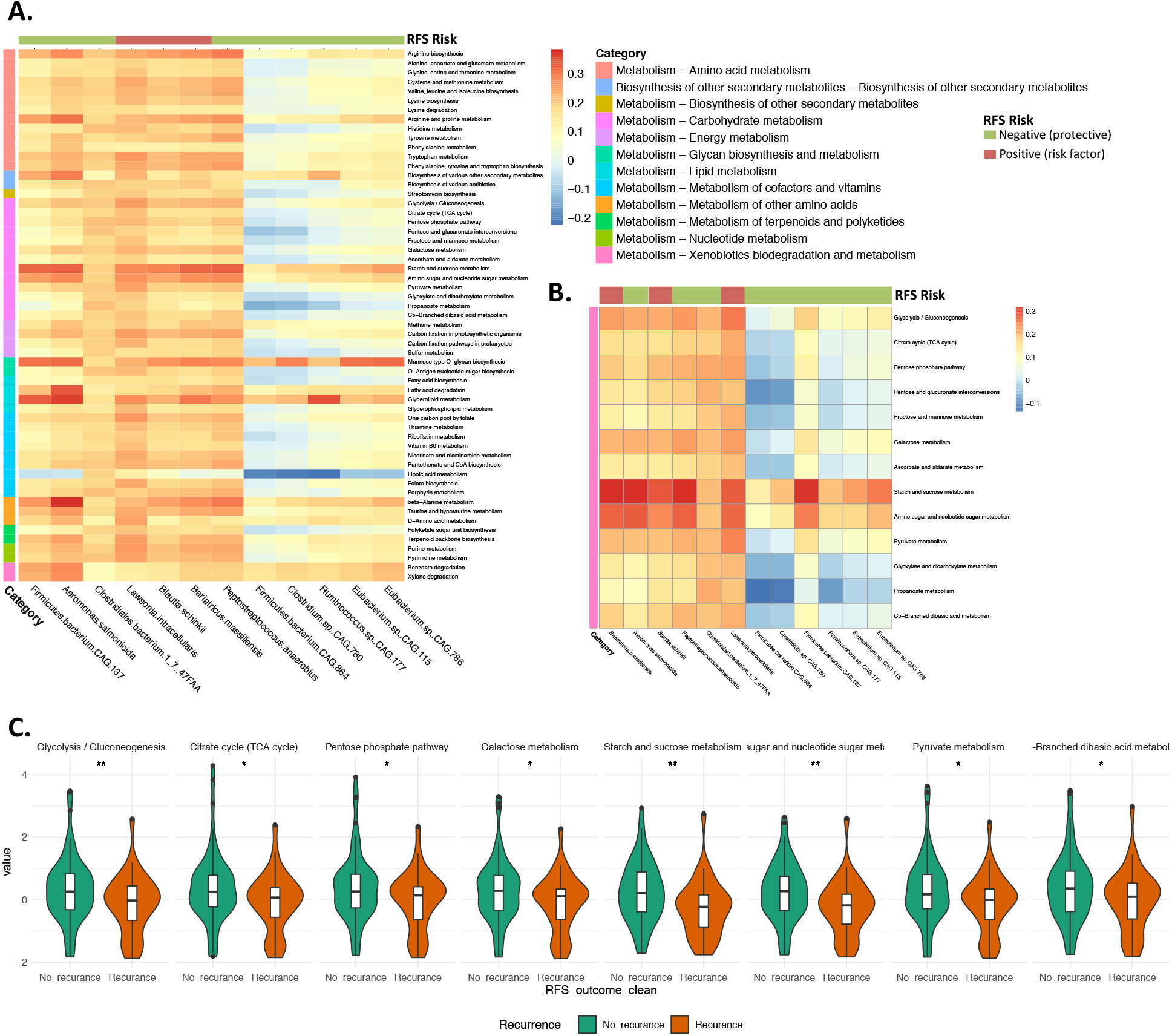
Association of Functional Pathways with Recurrence Biomarkers. (A) depicts the correlation between z-score normalized KEGG Level 3 bacterial pathways (presented in rows) of the recurrence-associated taxa (displayed in columns). The color of the column strip indicates the direction of association of the bacteria with recurrence (red for positive association and green for negative). The color of the row strip indicates the category (KEGG Level 2) of the functional pathway. The color of each cell represents the level of correlation. Functions are included only if they have a correlation FDR<0.0001 for a minimum of five recurrence-associated taxa. (B) exclusively displays Carbohydrate metabolism pathways derived from (A). (C) presents carbohydrate-associated pathways significantly correlated with recurrence in the North American region (no significant correlations in other regions), accounting for factors: age, sex, tumor stage, BRAF mutation and study arm. Y-axis of figure C represents the normalized z-score of the pathways.

### GMB matching facilitates cross-regional generalizability

Regional GMB heterogeneity is a major barrier to the development of reliable gut microbial markers for melanoma outcomes (He et al., 2018a). Recognizing this, we then tested whether recurrence-associated bacteria (**Figure 2A**) exhibited stronger prediction for recurrence in individuals selected for closely similar overall GMB composition (JSD distance), regardless of geographic region (**Figure 4A**). The prediction of recurrence in non-North American participants related to the North America-specific markers (**Figure 2A**) was strongest for those most closely matched to the North Americans on JSD distance (**Figure 4B**). Non-North American participants matching North American participants at JSD of ≤0.11 (n=61) showed an AUC of 0.88. Furthermore the AUC was highly inversely correlated to JSD similarity (correlation = –0.85, p<0.001), indicating that the smaller the beta-diversity distance between matched pairs, the stronger the prediction was in non-North American (validation set) participants of markers initially identified in North Americans (discovery set). Similar relationships were observed for markers initially identified in Western Europe (**Figure 4C**), Eastern Europe (**Figure 4D**), and ROW (rest of the world) (**Figure 4E**), with the strongest predictions among those most closely matched on JSD (e.g., JSD ≤ 0.11). Similar results are found for other measures of beta diversity (**Figure 4F**). While there were differences in number of patients retained using different beta-diversity measures (**Figure 4F**), the overall pattern was the same and consistently indicated that close GMB matching, regardless of distance metric choice, yielded more robust generalizability.

**Figure 4.**
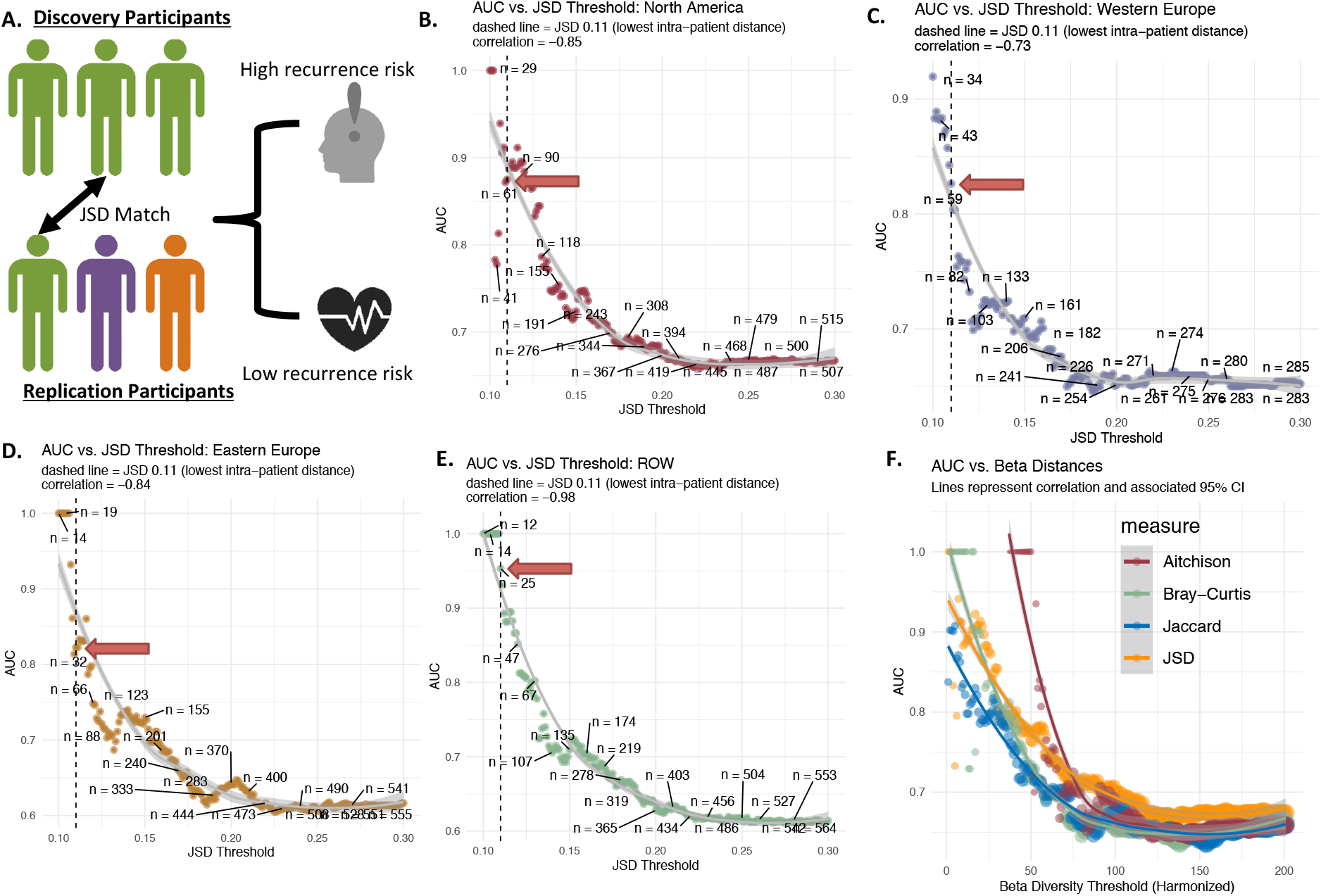
Recurrence Risk Prediction Models in Patients Using Independent Cross-regional Replicates Matched on GMB. **Panel A** depicts the patient matching method employed to generalize markers (i.e. using the region specific markers in other geographic areas for patients with the “same” GMB). JSD was used to match patients across region (testing patients are always from a different region from training patients), and subsequently, the predictive power of biomarkers was evaluated in the subsequent panels with adjustment for age, sex, tumor stage, BRAF mutation and study arm. **Panel B shows** the relationship between prediction measured using AUC vs. increasing JSD distance (spearman correlation = –0.85, p<0.001). For each point (200 total) a non-North American patient is matched to a North American subject at each JSD threshold and the final independently matched set is modeled using the North America markers to obtain AUC. **Panel C, D and E** show the same analysis, but using the ROW, Eastern Europe and Western Europe as discovery sets respectively. **Panel F** shows the comparison between the original JSD beta-diversity as well as Bray-Curtis dissimilarity, Jaccard index, and Aitchison dissimilarity with respect to AUC predictive power. Metrics were standardized by setting the lower limit to the median intra-sample distance and the upper limit to the median inter-sample distance, with 200 equal intervals for testing.

**Figure 5.**
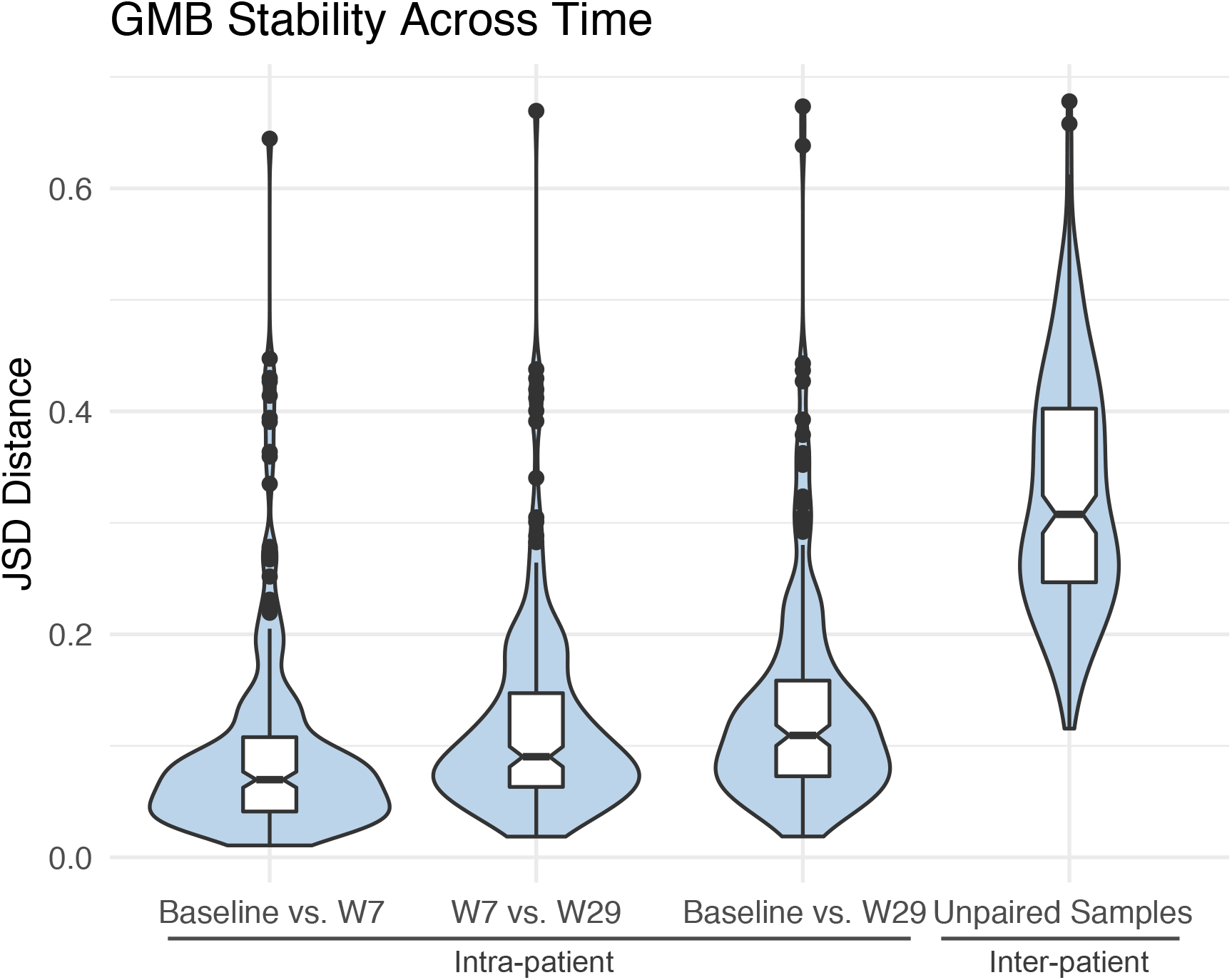
GMB Stability Across Time. Figure shows the bacterial β-diversity measured using Jensen Shannon divergence between measured visits (intra-patient variation) as well as between all unpaired samples for reference (inter-patient variation). Overall GMB was largely unchanged across baseline, week 7 and week 29 measurements with global PERMANOVA R^2^ = 0.867, p-val<0.001. Comparison between baseline and week 7 samples had an R^2^ = 0.930, p-val<0.001; week 7 vs. week 29 R^2^ = 0.900, p-val<0.001.

### Temporal Stability of GMB Following ICB Treatment

To assess the temporal stability of the GMB, we calculated intra-patient microbial JSD distances at baseline, week 7, and week 29, in 248 study participants with available serial stool samples, and as a comparison, we also calculated the unpaired inter-patient JSD distances (**Figure 4).** In this analysis, JSD values close to 0 indicate similarity, while JSD close to 1 indicate dissimilarity. We found that the GMB for individuals remained consistent across visits (global PERMANOVA across all three time-points, R^2^ = 0.867, p-value < 0.001), with remarkable stability of the GMB from before (baseline) and during (7 and 29 weeks) ICB therapy. The findings are consistent for oth treatment arms (nivolumab plus ipilimumab combination: R^2^ = 0.852, p<0.001 and nivolumab only R^2^ = 0.902, p<0.001) (see **Supplementary** Figure 2). Analysis of time-points as the outcome in place of patients did not reveal any significant compositional differences (**Supplemental Figure 4**), indicating that treatment did not have a targeted effect on the GMB (R^2^ = 0.0019, p-value=0.76). Longitudinal samples were more likely to be provided by those who didn’t experience recurrence during the trial (see **Supplemental Table 2**; recurrence rate: 27.2% for those providing longitudinal samples vs. 44.8% for those who didn’t). However, among those who provided samples, the GMB remained stable regardless of recurrence status (see **Supplement Figure 3**). GMB composition thus remained predominantly unchanged post-treatment without any identifiably consistent temporal shifts due to treatment, although a modest destabilization was noted for the combination treatment compared to the mono-treatment arm.

## Discussion

We investigated associations of the gut microbiome with melanoma recurrence, in the multi-center Checkmate 915 phase III trial of adjuvant immune checkpoint blockade (ICB), with nivolumab plus ipilimumab or nivolumab alone (Weber et al., 2023). We found that melanoma recurrence was associated with gut microbial taxa from the *Eubacterium, Ruminococcus, Firmicutes*, and *Clostridium* genera in region-specific and cross-region meta-analyses.

Recurrence prediction by these markers was best achieved across regions by matching on GMB compositional similarity between the intra-regional discovery and external validation sets. AUCs for prediction ranged from 0.83-0.94 (depending on the initial discovery region), for patients closely matched on GMB composition (e.g., JSD ≤0.11). This evidence indicates that taxonomic markers for prediction of recurrence are generalizable across regions, for individuals of similar GMB composition. Lastly, we examined longitudinal samples from patients during treatment and discovered that the GMB composition remained largely constant over the treatment period, indicating stability of the gut microbiome throughout the ICB treatment course.

We identified specific bacterial strains that predict recurrence in the adjuvant setting. The *Eubacterium*, *Ruminococcus*, *Firmicutes*, and *Clostridium* have been previously associated with outcomes for ICB in the metastatic setting (Lee et al., 2022; Matson et al., 2018; Routy et al., 2018). *Eubacterium* has been shown to modulate the efficacy of immunotherapies, by promoting an anti-inflammatory environment via natural killer cell interaction (Liu et al., 2023). Similarly, *Clostridium* and *Firmicutes* have been linked to enhanced immunoregulatory responses, related to modification of the T-cell response which may directly enhance the effects of immunotherapy (Shim et al., 2023). *Ruminococcus*, on the other hand, has been associated with both pro– and anti-inflammatory effects, particularly related to increases in CD4+ and CD8+ T cells, potentially influencing the outcomes of immune checkpoint therapies (Araji et al., 2022). Additional recurrence-associated taxa identified in this study include *Lawsonia, Bariatricus,* and *Blautia,* the latter of which was also associated with ICB treatment outcomes in our previous pilot research (Peters et al., 2019). The results shown here in the adjuvant setting and research in the metastatic setting (Lee et al., 2022; Matson et al., 2018; Routy et al., 2018) (Peters et al., 2019) are beginning to identify bacteria that impact ICB treatment outcomes, setting the stage for future studies modifying the GMB to achieve more favorable outcomes in a variety of ICB contexts.

Our analysis also showed connections between these immunomodulatory bacterial taxa and carbohydrate metabolism pathways within the GMB, with associations related to glucose metabolism being among the most numerous category. We observed significant correlations between certain bacterial taxa, such as *Eubacterium* and *Ruminococcus*, and the KEGG Level 3 pathways related to carbohydrate metabolism. This is notable because “glycolysis/gluconeogenesis” and “pentose phosphate pathway” have been related to both the microbiome and cancer treatment success (Cullin et al., 2021). Similar findings have been reported by Spencer et al (Spencer et al., 2021) who reported that dietary fiber can modulate the gut microbiome (specifically *Eubacterium* and *Ruminococcus*) and enhance the response to melanoma immunotherapy, implying that a high-fiber diet could shift the microbiome towards a composition conducive to enhanced immunotherapy response. Previous evidence (Spencer et al., 2021) along with our comprehensive study focusing on the adjuvant setting adds weight to the proposition that dietary interventions on the GMB are a potential strategy to reduce melanoma recurrence risk.

A barrier to progress in use of GMB biomarkers as tools for clinical prediction in ICB treatment of melanoma is that GMB markers associated with melanoma outcomes tend to be population specific (Lee et al., 2022), as geographic locality is a strong determinant of GMB composition (Peters et al., 2020). In our study, we also observed significant variation in GMB composition between regions internationally. We showed, however, that the capacity to predict recurrence may be improved by limiting comparisons to subjects closely matched for GMB beta-diversity. This implies, for practical application in the clinical setting, that prediction of recurrence for individual melanoma patients may be achievable by comparison to referent data for patients closely matched on GMB; this will require larger data sets well-characterized for GMB and ICB outcomes than are currently available.

In our study, an important design element included sampling of the GMB before and at several times during treatment, to assess ICB treatment-related changes in microbiome composition. While in free-living populations, GMB remains highly stable over a time course that may span years (Chen et al., 2021; Olsson et al., 2022), ours is the first study to demonstrate temporal stability of the GMB in ICB-treated patients. Given previously reported improvement in outcomes in ICB-treated melanoma patients by fecal microbiome transplant (FMT) (Derosa and Zitvogel, 2021), our results suggest that FMT or other GMB modifiers could exert a stable benefit throughout the treatment course. The stability of the GMB during ICB treatment, as illustrated in our data, hints at its potential as a lasting therapeutic reservoir which by alteration—whether through dietary changes, probiotics, or FMT—may offer a novel strategy for enhancing the effectiveness of adjuvant ICB treatment.

## Methods

### Study Population and Design

Our gut microbiome study was based on the phase III CheckMate 915 trial (ID: NCT02388906)(Weber et al., 2023), which originally evaluated adjuvant nivolumab plus ipilimumab versus nivolumab alone in patients with resected stage IIIB-D or IV melanoma. The primary endpoint was recurrence-free survival (RFS). The original trial reported that there was no significant difference between treatment groups for RFS. For a full description of original trial including outcome assessment and sample collection, refer to the original design publication (Weber et al., 2023). Our prospective, analysis focused on a total of 674 patients who provided a stool sample prior to treatment initiation (**Supplemental Figure 1**).

### Sample Collection and Sequencing

Participants had the option to provide stool samples prior to the commencement of their treatment. The ancillary microbiome study showed no significant differences compared to the original trial in clinical and demographic variables (Weber et al., 2023). Participants average age was 55 years. Most patients were stage IIIC, with a slightly higher proportion of men than women. In order to quantify the impact of treatment on GMB, during and after treatment, approximately half of the participants were also required to submit stool samples during their treatment (specifically at week 7) and post-treatment (at week 29). The stool samples have been collected using OMNIgene GUT kits (DNA Gentotek, Ontario, CA), which provide room temperature stability of microbiome profiles for 2 months. All samples have been collected during doctor visits and mailed by the patient to a centralized laboratory, per region, for storage where samples were immediately store at –80°C.

These samples underwent rigorous shotgun metagenomics sequencing in the Knight laboratory at the University of California San Diego (UCSD) as previously described (Usyk et al., 2023), enabling us to achieve strain-level resolution of the GMB. DNA was extracted from stool following the Earth Microbiome Project protocol (Thompson et al., 2017). Input DNA was quantified, using a PicoGreen fluorescence assay (ThermoFisher, Inc), and normalized to 1 ng, using an Echo 550 acoustic liquid-handling robot (Labcyte, Inc). Enzyme mixes for fragmentation, end repair and A-tailing, ligation, and PCR were added using a Mosquito HV micro pipetting robot (TTP Labtech). Fragmentation was performed at 37 °C for 10 min, followed by end-repair and A-tailing at 65 °C for 30 min. Sequencing adapters and barcode indices were added in two steps, following the iTru adapter protocol(Glenn et al., 2019). Universal “stub” adapter molecules and ligase mix were first added to the end-repaired DNA using the Mosquito HV robot and ligation performed at 20 °C for 1 h. Unligated adapters and adapter dimers were removed using AMPure XP magnetic beads and a BlueCat purification robot (BlueCat Bio). Next, adapter-ligated samples were added to a 384-PCR plate containing unique i7 and i5 index primers and PCR master mix, then PCR-amplified for 15 cycles. The amplified and indexed libraries were purified again using magnetic beads and the BlueCat robot, re-suspended in water, and transferred to a 384-well plate using the Mosquito HTS liquid-handling robot for library quantitation, sequencing, and storage. Samples were normalized and pooled based on a PicoGreen fluorescence assay, PCR cleaned, and size-selected on a PippinHT before sequencing on an Illumina NovaSeq 6000 (S4 flow cell and 2×150bp chemistry) at the Institute for Genomic Medicine at UCSD.

### Statistical and Bioinformatics Analysis

GMB composition was determined in samples by using the woltka pipeline with the wolR1 database executed on the Qiita platform (Gonzalez et al., 2018) using default pipeline settings. To discern patterns and significant differences in GMB, we employed global beta diversity analysis using Jensen Shannon divergence (JSD) (Fuglede and Topsoe, 2004) as the distance metric. JSD was selected because it was specifically determined to be highly effective for generalizability and direct utility in biomedical contexts (Sáez et al., 2017). Statistical significance of beta diversity measured using JSD was assessed by PERMANOVA (Anderson, 2014), using the vegan (Oksanen et al., 2007) package in R (Team, 2013). Specific strain/species markers were identified using ANCOM-BC (Lin and Peddada, 2020) for all outcomes (i.e. recurrence), with adjustment for age, sex, tumor stage, BRAF mutation and study arm.

Given the regional disparities in GMB compositions, a meta-analysis was deemed necessary. This was conducted on the identified region-specific markers to determine their overarching association with recurrence across all regions. Meta analysis was performed using the ANCOM-BC derived effect estimates and analyzed for pooled effect using the meta package in R using random effect meta-analysis (Schwarzer, 2007), employing the default settings (Van den Noortgate et al., 2013).

In the GMB matched analysis samples were selected on the basis of JSD similarity to participants from the each discovery region (always selecting replication to include samples outside of the discovery region). Specifically non-discovery region participants were checked for JSD distance against all participants from the discovery region starting at JSD 0.01 (i.e. high GMB similarity) and finishing at JSD 0.3 (i.e. low GMB similarity) with steps of 0.001. For a replication participant to enter an analysis at a given JSD threshold, they need to have a match to at least one discovery region patient with a JSD distance being equal to or lower than the defined threshold. For example at the 0.11 JSD threshold, roughly the level of patient to self distance, a participant would enter into the analysis if at least one discovery region subject has a JSD of 0.11 or lower to him (indicating high GMB similarity) and discarded if no such match could be made. Area under the curve analysis was performed using the pROC (Robin et al., 2014) package in R.

For the comparison between beta diversity metrics and recurrence prediction, we standardized metrics by setting the lower limit to the median intra-sample distance and the upper limit to the median inter-sample distance, with 200 equal intervals in between the bounds for testing AUC predicting on matched samples. This process of modeling for each involved selecting non-North American patients for testing using North American regional markers based on a beta-diversity threshold (each of the 200 steps), followed by AUC calculation in independent testing set.

### Outcome Measures

The primary outcome measure was the recurrence of melanoma post-treatment. Secondary outcomes included regional variations in these associations and stability of the GMB during the course of ICB treatment.

### Data Availability

The raw sequence data for all baseline samples along with uncontrolled phenotype variables reported in this paper have been deposited in the Genome Sequence Archive (Genomics, Proteomics & Bioinformatics 2021) in National Genomics Data Center (Nucleic Acids Res 2021), China National Center for Bioinformation / Beijing Institute of Genomics, Chinese Academy of Sciences (GSA: HRA005933, direct link: https://bigd.big.ac.cn/gsa-human/browse/HRA005933) that are publicly accessible at https://ngdc.cncb.ac.cn/gsa.

Additionally, all shotgun metagenomics sequencing data and the generated biom files containing bacterial taxa and functional profiles are available within Qiita under the StudyID 13059.

### Funding/Support

Research reported in this publication was supported in part by the U.S. National Cancer Institute under award numbers P50 CA225450, P20CA252728, P30CA016087, R01CA159036, R01LM014085-01A1, and U01CA250186.

## Acknowledgements

We thank all of the participants of the Checkmate 915 trial (ID: NCT02388906) for providing their samples and making this research possible. This publication includes data generated at the UC San Diego IGM Genomics Center utilizing an Illumina NovaSeq 6000 that was purchased with funding from a National Institutes of Health SIG grant (#S10 OD026929). Research reported in this publication was supported in part by the U.S. National Cancer Institute under award numbers P50 CA225450, P20CA252728, P30CA016087, R01CA159036, R01LM014085-01A1, and U01CA250186.

## Supplement Table of Contents

### Tables

Supplemental Table 1. Region Stratified ANCOM-BC Results

### Figures

Supplemental Figure 1. Study Consort Chart

Supplemental Figure 2. JSD Distances Across Time, Stratified by Trial Arm

Supplemental Figure 3. JSD Distances Across Time, Stratified by Recurrence Status

Supplemental Figure 4. PCOA Plot by Time Points

**Supplemental Table 1.** Region Stratified ANCOM-BC Results.

| Region | Recurrence Associated Strain | OR | 95% CI, low | 95% CI, high | p-val | q-val |
| --- | --- | --- | --- | --- | --- | --- |
| North America | Firmicutes bacterium CAG:137 | 0.17 | 0.25 | 0.11 | 1.64E-05 | 0.028 |
| North America | Firmicutes bacterium CAG:884 | 0.40 | 0.49 | 0.33 | 8.43E-06 | 0.015 |
| North America | Clostridium sp. CAG:780 | 0.21 | 0.30 | 0.15 | 9.26E-06 | 0.016 |
| North America | Eubacterium sp. CAG:115 | 0.054 | 0.10 | 0.030 | 9.76E-07 | 0.0017 |
| North America | Eubacterium sp. CAG:786 | 0.09 | 0.13 | 0.058 | 1.67E-10 | 2.89E-07 |
| North America | Peptostreptococcus anaerobius | 0.49 | 0.58 | 0.42 | 1.13E-05 | 0.020 |
| North America | Eubacterium siraeum | 0.088 | 0.15 | 0.052 | 3.39E-06 | 0.006 |
| North America | Ruminococcus sp. CAG:177 | 0.059 | 0.11 | 0.033 | 9.74E-07 | 0.0017 |
| North America | Aeromonas salmonicida | 0.42 | 0.51 | 0.34 | 2.37E-05 | 0.041 |
| Western Europe | Bariatricus massiliensis | 1.49 | 1.64 | 1.36 | 1.50E-05 | 0.028 |
| Western Europe | Blautia schinkii | 1.45 | 1.58 | 1.33 | 1.13E-05 | 0.021 |
| Eastern Europe | Lawsonia intracellularis | 2.50 | 3.09 | 2.02 | 1.62E-05 | 0.030 |
| Rest of World | Clostridiales bacterium 1_7_47FAA | 0.22 | 0.31 | 0.15 | 1.60E-05 | 0.032 |

Table shows the results of ANCOM-BC analysis of recurrence as the outcome with GMB as the core predictor with adjustment for participant age, sex, tumor stage, BRAF mutation and study arm. Odds ratios (ORs) are shown with associated p-values as well as adjusted values using the “holm” method.

**Supplemental Table 2:** Patients with Longitudinal Sampling.

| Variable | No | Yes | OR | pval |
| --- | --- | --- | --- | --- |
| N | 424 | 301 |  |  |
| Recurrence |  |  |  |  |
| No | 228 (55.2%) | 219 (72.8%) | 0.46 (0.33 - 0.64) | 1.68E-06 |
| Yes | 185 (44.8%) | 82 (27.2%) | - | - |
| Missing (n = 11) |  |  |  |  |
| Age Mean± SD | 54.54± 14.32 | 55.4± 13.3 | - | 0.685 |
| Gender |  |  |  |  |
| Female | 178 (42.4%) | 120 (39.9%) | 1.11 (0.81 - 1.52) | 0.54 |
| Male | 242 (57.6%) | 181 (60.1%) | - | - |
| Missing (n = 4) |  |  |  |  |
| Region |  |  |  |  |
| Australia | 77 (18.2%) | 64 (21.3%) | - | 0.13 |
| Eastern Europe | 24 (5.7%) | 21 (7%) | - | 0.125 |
| ROW | 29 (6.8%) | 19 (6.3%) | - | 0.284 |
| North America | 66 (15.6%) | 39 (13%) | - | 0.295 |
| Western Europe | 224 (52.8%) | 158 (52.5%) | - | 0.148 |
| Stage at entry |  |  |  |  |
| Not Reported | 7 (1.7%) | 0 (0%) | - | 1 |
| Stage IIIB | 113 (26.7%) | 102 (33.9%) | - | 0.125 |
| Stage IIIC | 234 (55.2%) | 152 (50.5%) | - | 0.16 |
| Stage IIID | 10 (2.4%) | 5 (1.7%) | - | 0.53 |
| Stage IV | 56 (13.2%) | 42 (14%) | - | 0.141 |
| B.Raf.Mut |  |  |  |  |
| Invalid/Not Reported | 104 (24.5%) | 75 (24.9%) | - | 0.145 |
| Mutant | 119 (28.1%) | 79 (26.2%) | - | 0.157 |
| Wildtype | 197 (46.5%) | 147 (48.8%) | - | 0.141 |
| Melanoma Subtypes |  |  |  |  |
| Acral | 14 (3.3%) | 9 (3%) | - | 0.268 |
| Cutaneous | 357 (84.2%) | 264 (87.7%) | - | 0.142 |
| Mucosal | 4 (0.9%) | 1 (0.3%) | - | 1 |
| Not Reported | 8 (1.9%) | 0 (0%) | - | 1 |
| Other | 37 (8.7%) | 27 (9%) | - | 0.146 |
| LD_baseline Mean± SD | 218.92± 85.25 | 213.48± 88.16 | - | 0.207 |

**Supplemental Figure 1.**
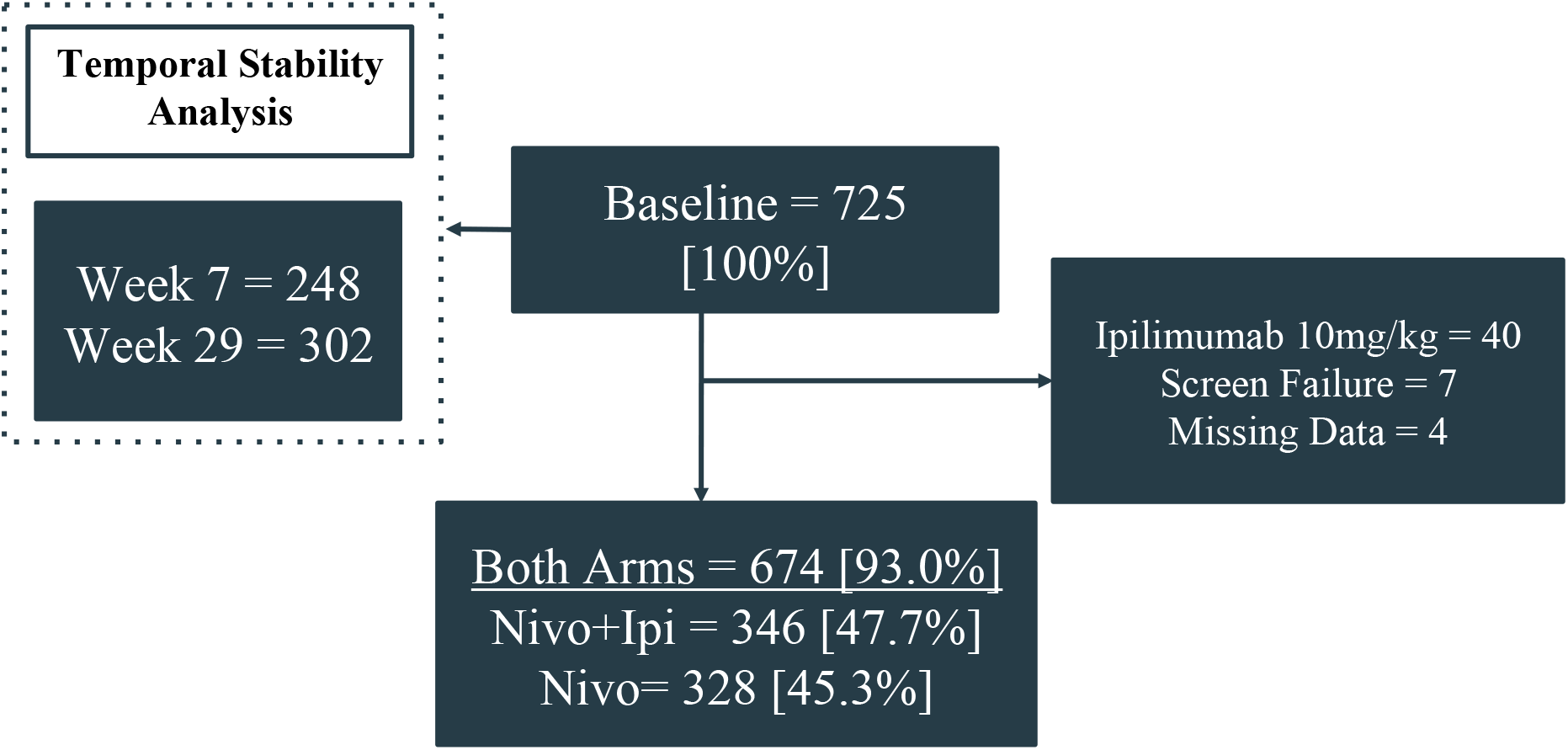
Study Consort Chart. 725 represented baseline. Of these, approximately half of the patients had follow-up sampling at weeks 7 and 29. From the baseline samples, 51 individuals were excluded due to coming from a supplementary arm of the original trial (n = 40), being screen failures (n = 7) or having missing randomization data (n = 4). Overall we utilized 674/725 (93.0%) of the available shotgun metagenomic samples for our core analysis.

**Supplemental Figure 2.**
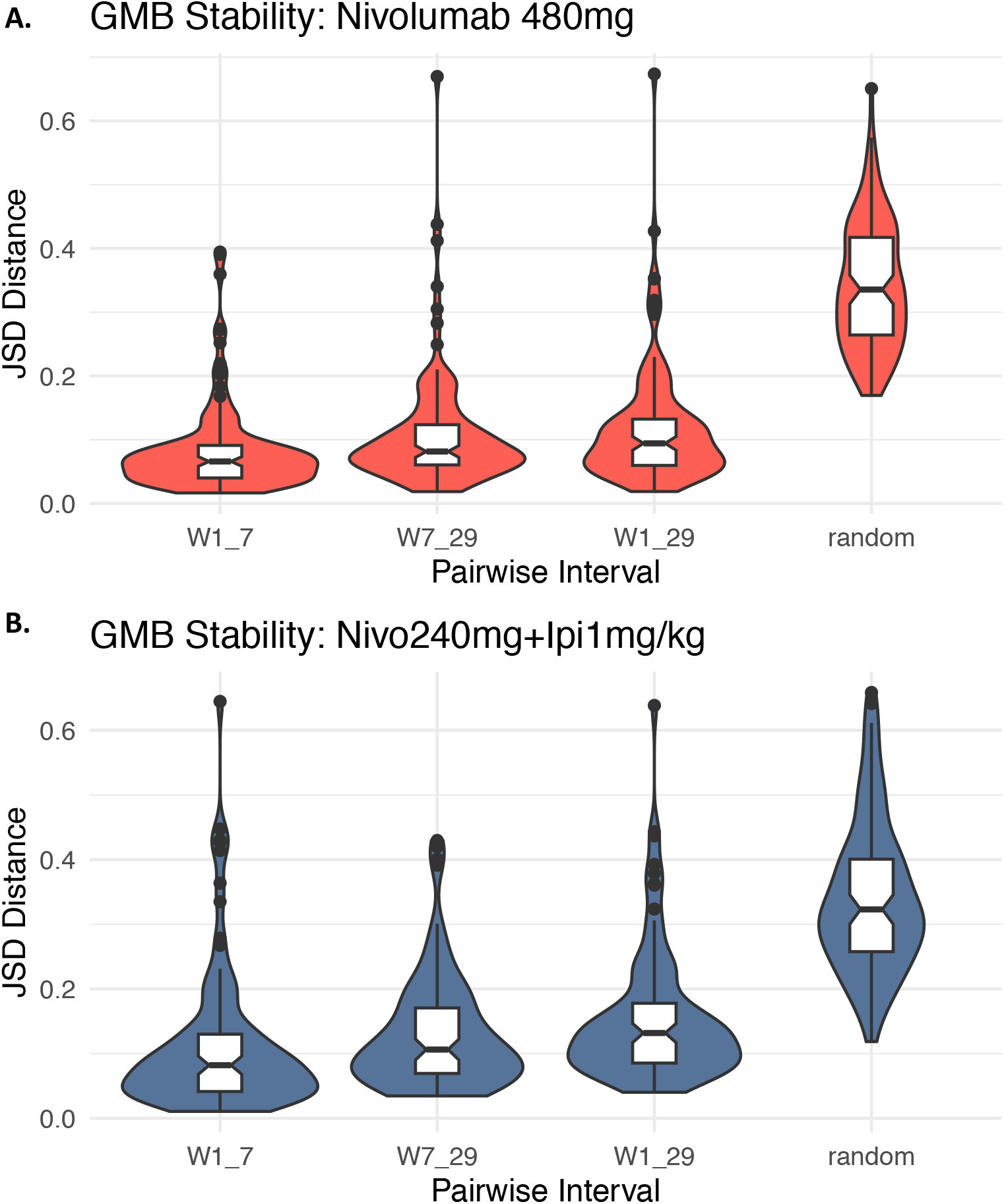
JSD Distances Across Time, Stratified by Trial Arm. The figure shows the bacterial β-diversity measured using Jensen Shannon divergence between measured visits (intra-patient variation) as well as between all unpaired samples for reference (inter-patient variation), stratified by the treatment arm (red, panel A is mono treatment and blue panel B is combination treatment). Overall GMB was largely unchanged across baseline, week 7 and week 29 measurements in both arms.

**Supplemental Figure 3.**
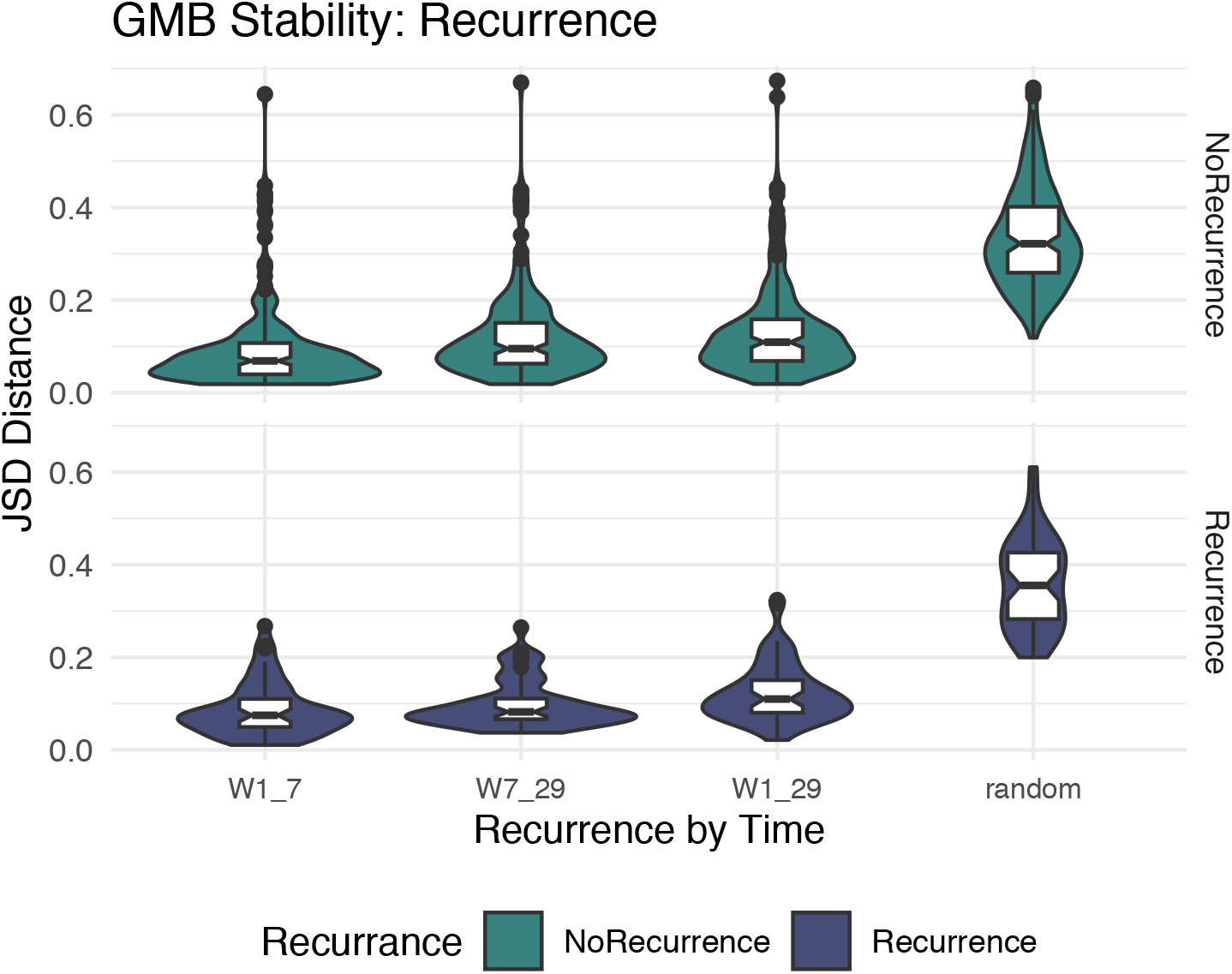
JSD Distances Across Time, Stratified by Recurrence Status. The figure shows the bacterial β-diversity measured using Jensen Shannon divergence between measured visits (intra-patient variation) as well as between all unpaired samples for reference (inter-patient variation), stratified by recurrence status (green, panel A: no recurrence group and navy, panel B, recurrence group). Overall GMB was largely unchanged across baseline, week 7 and week 29 measurements in both panels.

**Supplemental Figure 3.**
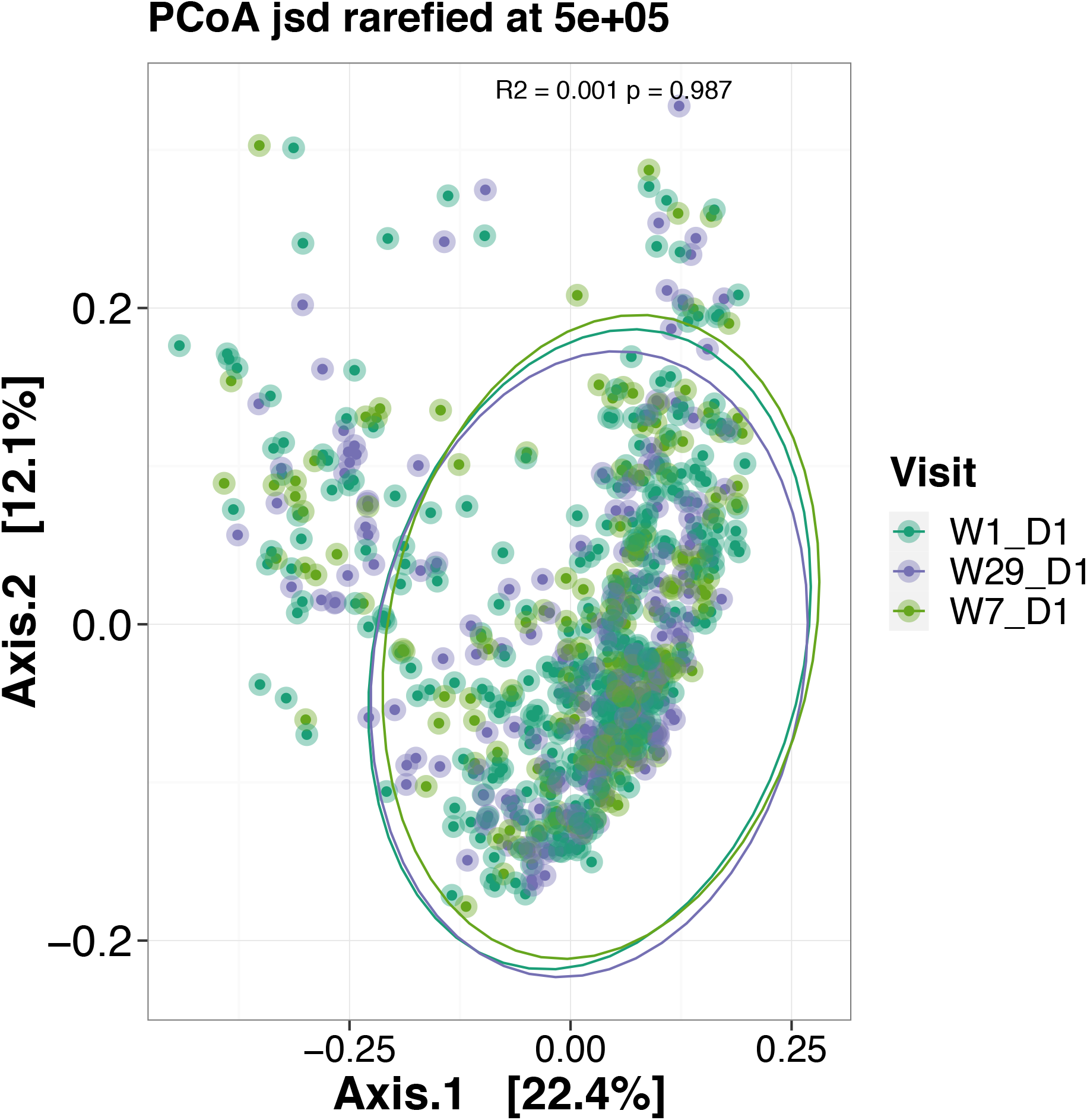
PCOA plot by Time Points. PCOA plat for three-time points as the outcome for the BMS patients using JSD distances.

